# Limitations of the refolding pipeline for *de novo* protein design

**DOI:** 10.64898/2025.12.09.693122

**Authors:** Kerlen T. Korbeld, Vsevolod Viliuga, Maximilian Fürst

**Affiliations:** Molecular Enzymology Group, University of Groningen, Nijenborgh 3, Groningen, 9747AG, The Netherlands; Department of Biochemistry and Biophysics and Science for Life Laboratory, Stockholm University, 171 21 Solna, Sweden; Max Planck Institute for Polymer Research, Ackermannweg 10, 55128 Mainz, Germany

**Author notes:** Equal contribution (Kerlen T. Korbeld, Vsevolod Viliuga).

## Abstract

With the emergence of powerful deep learning-based tools, computational protein design has become a widely accessible technique. Nowadays, it is possible to perform both sequence and structure design in a matter of minutes, making the technology attractive to the broader scientific community. In protein design campaigns, one of the most common *in silico* strategies to evaluate how well a sequence encodes a target structure is the so-called self-consistency or refolding pipeline. In this approach, a structure prediction model is used to refold the designed sequence to probe whether it is compatible with the intended structure, and is evaluated via two metrics linked to experimental success: the confidence score of the predicted structure (pLDDT) and the self-consistency root-mean-square deviation (scRMSD), which measures how closely the refolded structure matches the target. In this work, we systematically evaluate how different models and structure prediction settings impact these metrics, and to what extent they can be used to reliably filter sequence design candidates. We show that evolutionary information can obscure folding models’ abilities to assess sequence-structure compatibility, reducing the predictive performance of refolding metrics for experimental success, particularly for designs that share homology with natural sequences. We further highlight limitations of refolding metrics, including their sensitivity to structural features, such as flexibility. Our findings raise awareness of potential pitfalls in refolding-based evaluation and support more informed use of these metrics in protein design campaigns.

## 1 Introduction

The emergence of highly precise protein structure prediction methods, such as AlphaFold2 (AF2)^1^ and ESMFold^2^ has catalyzed a paradigm change in the field of protein design. Enabled by these and other breakthroughs in deep learning (DL), *de novo* design has rapidly expanded from a niche technology with modest, albeit striking achievements, to a now widely applicable method.^3^ Building on the DL architectures for protein structure prediction, many new computationally efficient and increasingly user-friendly methods for both backbone and sequence design complement established physics-based approaches like Rosetta.^4,5^ Strikingly, the field is seeing dramatic improvements in experimental success rates, with designs that successfully express in cellular hosts now often displaying remarkable structural similarity to computational models, highlighting the accuracy of state-of-the-art modelling tools.^6,7^ These rapidly advancing algorithmic and technical developments have enabled notable achievements, including the design of multi-chain complexes with novel topologies,^8^ highly affine binders for therapeutically relevant protein targets,^9,10^ solubility-optimized redesign of enzymes with improved catalytic efficiency,^11^ and even shape-conditioned structures encoding letters from the English alphabet.^12,13^ Given these advances, DL-based protein design matured into a reliable tool used both in fundamental as well as applied biological research.

Protein design studies are almost always driven by the goal of achieving a particular biological function, and thus usually comprise identifying a function-compatible protein structure, as well as an amino acid sequence that encodes that structure.^14,15^ A broad distinction is made between redesign tasks, in which the functional or physicochemical properties of a natural template are optimized, and *de novo* design, employed for instance when no functional equivalent in biology is known. Depending on the goal and strategy, various tools may be applied for *de novo* design. The most established design pipeline is, however, a two-step approach: first, a protein backbone is generated *de novo*, and a second model predicts an amino acid sequence encoding the target structure in a downstream step (Fig. 1). Numerous specialized and general-purpose tools have been developed for backbone generation, with diffusion-based generative models such as RFdiffusion^8^ currently exhibiting a particular prominence. In contrast, fewer options exist for sequence design, which is currently dominated by the ProteinMPNN family of models.^16,17^ While models that predict a sequence given structure are often referred to as inverse folding models, the term was originally defined more strictly as identifying a sequence that solely adopts the target, and no other low-energy conformation.^18,19^ As this is ultimately the desired outcome, most *in silico* pipelines include an evaluation step that probes the absence of these off-target folds for designed sequences.

**Figure 1.**
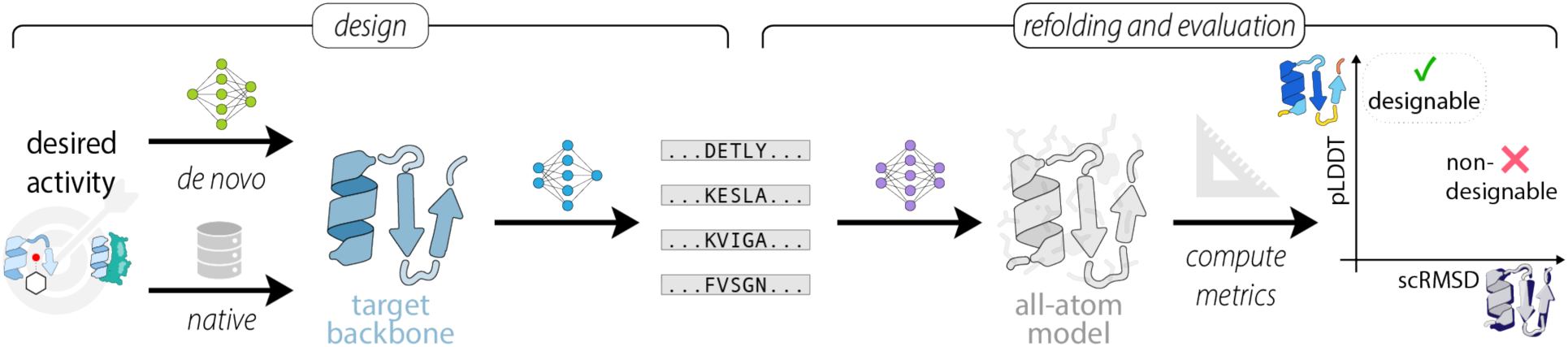
Designability assessment with the refolding pipeline in a fixed-backbone design task. Starting from a target structure of a native or a *de novo*-designed protein, a sequence design model generates a pool of candidate sequences. Next, structures of the designed sequences are refolded using a structure prediction model and are subsequently superposed onto the target structure. If the predicted structure is of high confidence (high pLDDT) and exhibits little deviation from the target in Euclidean space (low scRMSD), the design is considered successful.

Notably, despite a diversity in design objectives and approaches, the majority of design campaigns rely on a very similar strategy for this evaluation. The most widely implemented approach in the field involves subjecting the designed sequence to a structure prediction model, i.e. the reverse of the sequence design process (Fig. 1). The basis for using refolding in this way lies in the assumption that folding models have implicitly learned an approximation of the energetics of protein structures and thus can be used to assess sequence-structure compatibility.^20^ In its entirety, this so-called refolding or self-consistency (sc) pipeline assesses whether a designed sequence reliably encodes the target structure. Importantly, structure prediction serves not only as an orthogonal quality control step on the output of the design workflow, but is also widely used as a quantitative tool to filter and rank sequence candidates. Design quality is generally assessed using two metrics derived from the predicted model: the predicted local distance difference test (pLDDT), and the structural deviation from the design target, usually measured as the self-consistency root mean square deviation (scRMSD) computed by superimposing the predicted structure onto the target backbone. Only sequences with pLDDT values above and scRMSDs below a predefined threshold are chosen for further characterization.^8,16,21^ Following a recent terminology shift, which extends a concept initially reserved for backbones,^22^ such selected sequences may be referred to as “designable” (Fig. 1).^23,24^

Notably, no standardized benchmarks or reference values for the scRMSD and pLDDT thresholds exist. Rather, each study inconsistently chooses the values *ad hoc*. While often, the scRMSD is computed via a global C*α*-atom superposition with a designability threshold set to 1.5-2 Å,^23^ some studies use the structure alignment’s template modeling (TM) score, with a common threshold of 0.5.^21,25,26^ pLDDT thresholds show even less consensus, ranging from 70 to 90 across studies.^11,16,24,27^ Moreover, despite the widespread use of the refolding pipeline,^16,24,27^ there remains an uncertainty over the degree of correlation between designability criteria and experimental success. A compounding issue is that both scRMSD and pLDDT can be critically influenced by the folding model and the choice of settings for the structure prediction step, which also differ. Protein design campaigns focused on the redesign of protein sequences frequently use AF2 with multiple sequence alignments (MSAs), either provided explicitly^11^ or generated automatically by ColabFold^28^ in its default settings.^29,30,31,32^ In contrast, most studies focusing on *de novo* structure generation followed by sequence design report depleting AF2 of evolutionary information by running predictions in single sequence mode,^24,25,27,33^ or use the ESMfold structure prediction model.^13,34,35^

As there has never been a dedicated evaluation on how applying designability thresholds affects experimental success rates—widely referred to as the ability of structure prediction to act as an “oracle”^36^—we resolved to attempt a systematic analysis of available data. We hypothesized that the inconsistent use of models and metrics may introduce reproducibility issues at best and systematic errors at worst, thereby undermining the reliability of the refolding pipeline and promoting the selection of suboptimal designs for experimental testing. In this work, we thus investigate how different prediction and evaluation settings affect *in silico* designability in the context of a fixed backbone design task. We first demonstrate that for both natural and *de novo* designed sequences, the presence of evolutionary information in the structure prediction step reduces the folding models’ ability to assess sequence-structure compatibility, thereby reducing the predictive power of refolding metrics for experimental success. We next investigate the limitations of running the refolding pipeline without evolutionary information and highlight the inherent limitations of refolding-based designability. Specifically, we show that certain structural features, such as disordered or flexible regions, can confound refolding metrics, leading to less reliable designability assessments.

## 2 Results

### 2.1 Evolutionary information impairs the discriminative ability of the refolding pipeline

We began our investigation by asking how well folding models used in a classical refolding pipeline distinguish good from bad sequence designs, that is, those that fold into the desired structure and those that do not. Concomitantly, we assessed how this discriminative ability is affected by the inclusion of evolutionary information, which is provided explicitly to structure prediction models like AF2 in the form of an MSA, or implicitly in the form of ESM embeddings.^37^ We hypothesized that if a sequence design shares significant identity with the natural template, both information-rich sequence alignments as well as protein language model embeddings could strongly bias the folding model towards native-like structure predictions. As a result, even badly designed sequences might be assigned inflated, artificially high confidence metrics, compromising the reliability of the designability assessment.

To test this, we built a dataset composed of small, structurally distinct natural proteins from the SCOPe database (Methods 4.3-4.4), for which abundant evolutionary information, i.e. a deep MSA is present (Fig. S1, SCOPe WT). Although AF2 normally relies on MSAs,^20^ it can also predict structures from single sequences (AF2 ss). To enable a fair comparison between designability computed with and without MSAs, we trimmed the set to 50 randomly selected proteins that are classified as designable when predicted by AF2 ss using a lenient pLDDT threshold of 70 (Methods 4.2). Starting from their native sequences, we then gradually mutated 10% to 60% of arbitrarily selected residues to random amino acid tokens, thereby simulating badly designed sequences with variable identity to the native sequence. For each fraction of mutated residues, we generated 64 “designs” to reduce the impact of sequences that by chance may contain less disruptive mutations or share similarities to natural homologs. We then ran structure predictions for all mutated sequences using different models and input settings. In addition to ESMfold with default settings, we used AF2 ss or AF2 provided with a precomputed MSA (AF2 MSA) (Methods 4.2). Additionally, we reasoned that we could assess the effect of the native MSA context on the model’s confidence metrics by predicting structures with the MSA signal at mutated positions omitted. We thus ran another set of AF2 predictions using MSAs in which entire alignment columns were masked at substitution sites (AF2 MSA mask). For all approaches, we computed designability as the fraction of predicted structures satisfying scRMSD ≤ 2.0 Å and a 70, 80, or 85 pLDDT threshold (Methods 4.1, Fig. 2a).

**Figure 2.**
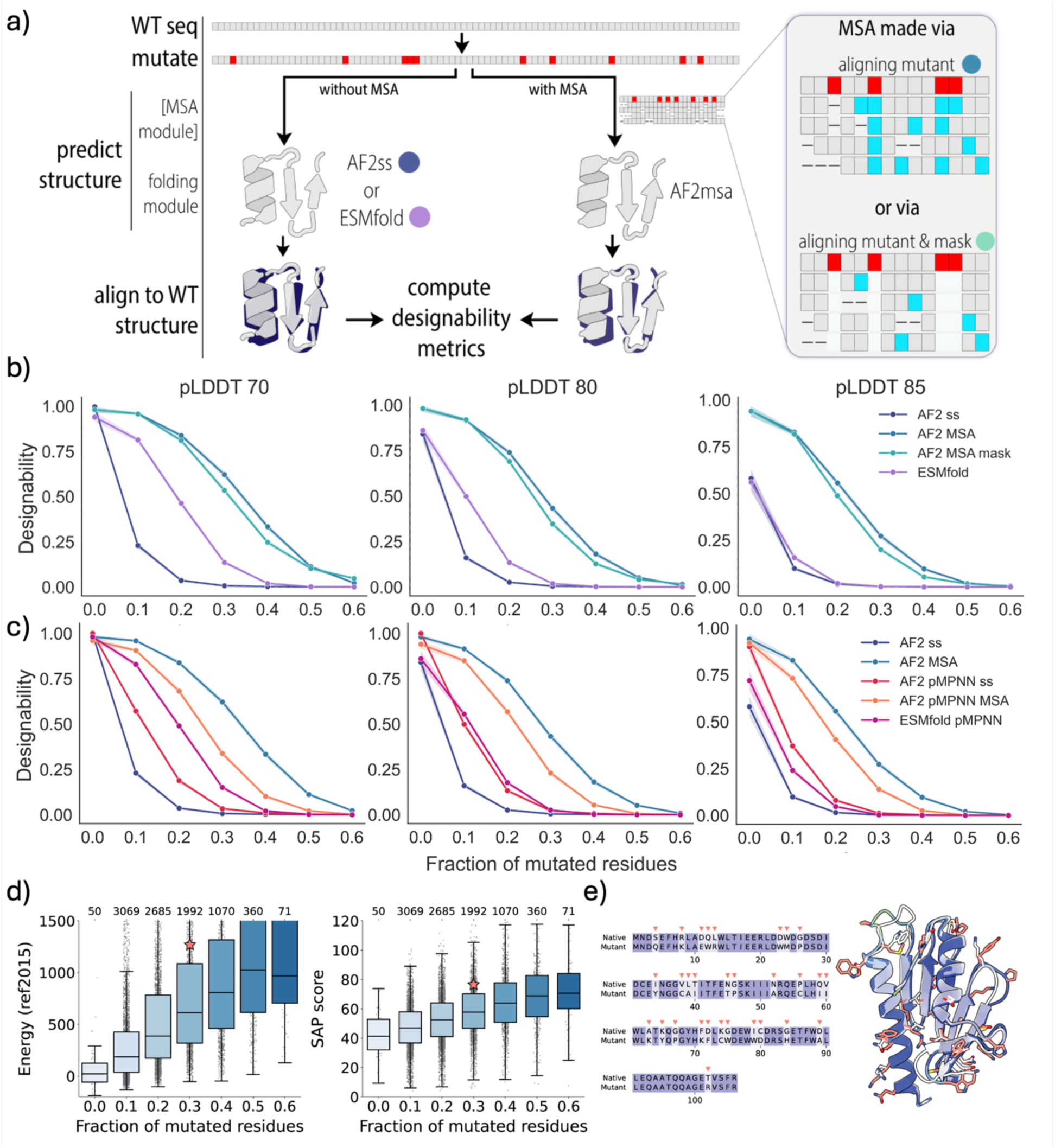
**a)** Schematic overview of the mutagenesis experiment. Random positions in native sequences were randomly mutated at specified percentages and given to MSA-free structure predictors like AF2 in single sequence mode or ESMfold. Alternatively, AF2 was given MSAs of homologous sequences to the mutant (AF2 MSA) or the same alignment masked at mutated positions (AF2 MSA mask). b) Designability of 50 random SCOPe proteins at various mutagenesis percentages. c) Same as in b) but using pMPNN designs as a starting sequence. d) Rosetta energy and spatial aggregation propensity (SAP) scores for structures classified as designable by AF2 MSA in b); sample counts indicated by numbers on top. e) Sequence and structure alignment of an example of a high-confidence structure of a mutant deemed designable by AF2 MSA at 30% mutated residues (SCOPe: d1ew4a, pLDDT 94, scRMSD 1.3). Its energy and SAP values are indicated by stars in d) and e). Mutant in salmon, native in white.

If folding models were reliable “oracles” of poor designs, we would expect a sharp decrease in designability upon random mutagenesis, given that introducing more than a handful of random mutations into natural proteins quickly leads to misfolding.^38^ Strikingly, however, we found that AF2 with explicit MSAs predicted structures of strongly mutated sequences with high confidence and low scRMSD to the native target. At 10% mutations, nearly all sequences are classified as designable by AF2 MSA regardless of the pLDDT threshold applied (Fig. 2b). Even at 30% mutations, over 60% of mutants remain designable with a pLDDT ≥70, while stricter thresholds of 80 and 85 reduce the designable fraction to about 40% and 30%, respectively. The AF2 MSA mask strategy produced nearly identical results, suggesting that even partial evolutionary information in otherwise deep MSAs is sufficient to bias AF2 towards making high-confidence predictions. Contrarily, AF2 ss demonstrated a faster drop in the mutants’ designability: at 20% mutations and higher, almost no sequences were classified designable. ESMfold also performed substantially better than AF2 MSA, and at a pLDDT ≥85 designability threshold approached the performance of AF2 ss. However, since AF2 ss and ESMfold rarely predicted the target structures of the unmutated natural sequences with pLDDTs above 80, the overall designability became relatively low at these thresholds (Fig. 2b).

In these experiments, many natural sequences (without mutations) predicted with ESMfold and AF2 ss already yielded structures with metrics below the designability thresholds. To create a dataset with higher fractions of designable sequences, we redesigned the SCOPe proteins with ProteinMPNN (pMPNN) (Methods 4.4), a sequence design model known to yield higher pLDDT structures in single sequence mode.^16^ For this dataset, we again mutated at defined fractions and computed designability as described for the native sequences. Crucially, we found that the discriminative power of AF2 ss decreased for mutants in the pMPNN background, compared to natural ones. At 10% mutated positions, the fraction of designable sequences increased from about 10% to nearly 60%, and at 20% mutations from nearly 0% to 20% at a pLDDT threshold of 70 (Fig. 2c). Indeed, this shift can be attributed to the overall higher pLDDT values assigned by AF2 ss to the pMPNN sequences (Fig. S2). In contrast to AF2 ss, ESMfold’s performance remained nearly unchanged compared to the experiment with mutants derived from natural sequences. While AF2 MSA’s discriminative power improved marginally for pMPNN-derived mutants, it remains very poor. Interestingly, although pMPNN shows about 41% native sequence recovery in our dataset, the MSAs computed for pMPNN-derived mutants remain nearly as deep as those of natural sequences (Fig. S2), which appears sufficient to bias structure predictions towards the natural homologs.

To address the possibility that the folding models’ high score for the randomized sequences might be due to a high fraction of them still being able to fold into the target, we computed energies and spatial aggregation propensities (SAP) with Rosetta^4,39^ for all mutants deemed designable by AF2 MSA. We observed a clear trend of markedly elevated median energies and SAP scores with an increasing fraction of mutated residues (Fig. 2d), strongly suggesting their unphysical nature. Visual inspection of highly mutated variants that yielded predicted structures with high pLDDT and low scRMSD confirmed this notion; the structures frequently showed multiple hydrophobicity and charge-altering mutations on both surface and in the core, highly likely to disrupt structural integrity of the fold (Fig. 2e).

These experiments suggest that refolding metrics should be treated cautiously when evolutionary information is included, where folding models struggle to flag designs with inappropriate residues even at high frequencies. While discriminative power increases if MSAs are absent, overall performance drops, thereby limiting the ability of folding models to assess sequence-structure compatibility of designed sequences.

### 2.2 Evolutionary information reduces the predictive power of refolding metrics for experimental success

After observing that providing evolutionary information reduces the ability of folding models to discriminate between nonsense designs and natural or pMPNN-designed sequences encoding natural backbones (Fig. 2), we next sought to examine this behavior in real-life protein design datasets. Using literature data from diverse protein design studies reporting experimentally tested sequences across different design pipelines and protein topologies (Fig. 3a) (Methods 4.5), we assessed the folding models’ ability to distinguish successful designs from failures. Notably, such analyses are complicated by factors that blur the distinction between truly bad designs—where a physical inability to fold prevents expression—from those merely affected by experimental factors—e.g. unfavorable transcription or protein degradation due to serendipitous cellular binding. We reasoned that if folding models were unaffected by the latter, such effects should occur randomly across all sequences and thus not systematically skew the models’ abilities to act as oracles in one or the other direction.

**Figure 3.**
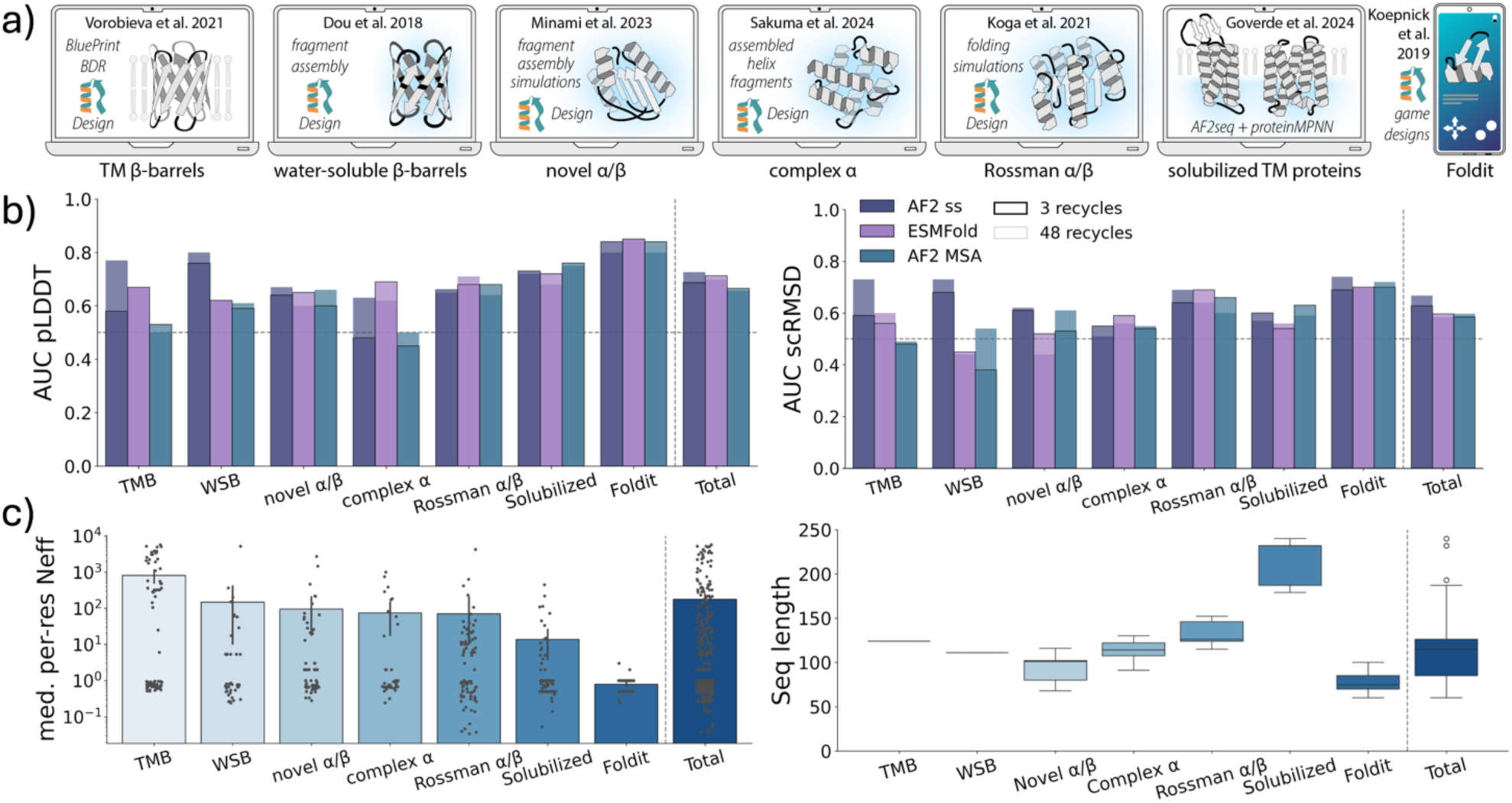
**a)** Datasets used in this experiment. Rosetta-based workflows are indicated with the Rosetta logo. b) Area under the ROC curve (AUC) for pLDDT and scRMSD as predictors of experimental success across various literature protein (re)design datasets for either the default 3 recycles (black outline) or 48 recycles (no outline). The horizontal line marks AUC 0.5 where predictive power is random. c) Sequence length distributions and MSA depths, expressed as median per-residue Neff. Neff values were calculated by counting non-gap residues at each MSA position using a weighting scheme and taking the median.

To test whether this was the case and rule out that the folding models’ training led it to in some way internalize experimental expression success, we tested whether models could distinguish between natural (and thus certainly physically plausible) proteins that do or do not express heterologously. To that end, we made use of the SoluProt dataset, which contains binary expression data for thousands of natural proteins.^40^ To be able to calculate scRMSDs and compute designabilities, we selected all proteins for which experimental structures are available, and further selected subsets of the soluble and insoluble entries with identical length distributions (Methods 4.5, Fig. S3). In addition, we performed pLDDT-only calculations for 1000 entirely randomly selected SoluProt sequences. As expected, none of the folding models showed discernible ability to predict heterologous protein expression success: the designable fraction did not differ significantly between soluble and insoluble proteins with available structures (Fig. S4), and the median pLDDTs of the unbiased set were also identical (Fig. S5). We therefore conclude that any discriminatory power we would observe in our analysis would largely be attributable to the model’s assessment of physical stability, rather than an implicit “knowledge” about experimental effects.

We then proceeded to predict structures of the literature-collected designs with AF2 MSA, AF2 ss, or ESMfold and probed their ability to identify successful designs. We computed pLDDT and scRMSD metrics for all 493 sequences and assessed whether they passed designability thresholds. Throughout the text, we refer to the experimental success if 1) the designed sequence could be expressed, 2) remained soluble, and 3) folded into the desired state, as verified by monodispersity in SEC or through exhibiting expected CD spectroscopy profiles. To facilitate comparison across studies that used various designability thresholds in their analyses, we computed threshold-independent areas under the curve (AUC) of receiver operating characteristic (ROC) curves.

While evolutionary information had negligible impact in some cases, it significantly impaired the predictive power of pLDDT and scRMSD in several datasets (Fig. 3b). The strongest performance drop occurred for the two β-barrels datasets, while a small difference was observed for the novel α/β and complex α sets. When investigating the cause of the variability across the datasets, we found no apparent correlation with sequence length, but noted that it was likely driven by the underlying MSA depths (Fig. 3c): since the sequences in two datasets (the Solubilized and Foldit datasets) had no or very few homologous sequences in databases, the resulting empty or very shallow MSAs were essentially equivalent as input for AF2 to single sequence mode. Interestingly, while another study reported ESMfold to outperform AF2 MSA as an oracle,^41^ our analysis suggests it remains inferior to AF ss except for datasets that fall into a shallow MSAs regime. As previously noted,^24^ we found that optimal folding of the transmembrane β-barrels required running folding models with more recycles. Overall, however, running models with few or many recycles did not substantially influence the results (Fig. 3b).

To better understand AF2 MSA’s poorer performance, we computed the difference in precision and F1 scores between AF2 ss and AF2 MSA at varying scRMSD and pLDDT cutoff combinations. For the deep MSA datasets, we found that the highest precision and F1 scores are consistently achieved with AF2 ss, confirming it as the better choice for reasonable thresholds (Fig. S6a). This effect is largely driven by a lower false positive rate for AF2 ss (Fig. S7), mirroring our previously noted tendency of AF2 MSA to be overly confident about poor designs. However, single sequence mode is not uniformly superior (Fig. S8): depending on dataset, MSA mode can match or exceed its performance at specific threshold combinations, and the location of the optimum varies widely (pLDDT 70-90; scRMSD 1-4 Å).

Although our identification of these optima is inherently retrospective and cannot prescribe thresholds for a new design campaign, the results indicate that design campaigns could benefit from adjusting thresholds after an initial experimental round. For the first pass, the practical reality of choosing thresholds to achieve a reasonable class balance has currently no good alternative. Importantly, our findings show that evolutionary information can directly alter refolding metrics and, in some cases, diminish their reliability as predictors of experimental success.

### 2.3 Limitations of AF2 in single sequence mode for refolding-based designability assessment

So far, our analysis has established that, despite limited overall performance, AF2 ss best identifies unphysical sequences as undesignable (Fig. 2) and most frequently predicts experimental success correctly across our datasets (Fig. 3). We next sought to investigate practical limitations of its applicability and potential remedies. Previous studies^2,20^ and our mutagenesis experiment indicated that AF2 struggles to predict accurate structures of natural sequences in single sequence mode, whereas pMPNN or protein language model designs yield high confidence models.^16,25,42^ However, whether prediction settings or specific structural features could impair AF2’s applicability for designability assessment has not been thoroughly examined.

To investigate this at scale, we computed the designability of a cleaned subset of the SCOPe dataset consisting of approximately ten thousand domains (Methods 4.6), using both the natural sequences and pMPNN designs with the lowest scRMSD to the natural target structure. We first assessed the influence of protein length on metrics computed with AF2 ss and found a strong inverse correlation: while already only 40% of the shortest native sequences in our test (up to 50 amino acids) passed the permissive designability thresholds of pLDDT ≥70 and scRMSD ≤2.0 Å, this fraction quickly approached 0 beyond a length of 150 residues (Fig. 4a-c). In contrast, pMPNN designs exhibited a substantial increase in designability across all lengths. While natural sequences rarely reached pLDDT values above 70, pMPNN designs consistently exceeded this threshold up to 200 residues and achieved average scRMSD values near 2 Å (Fig. 4a-c) up to a length of 150.

**Figure 4.**
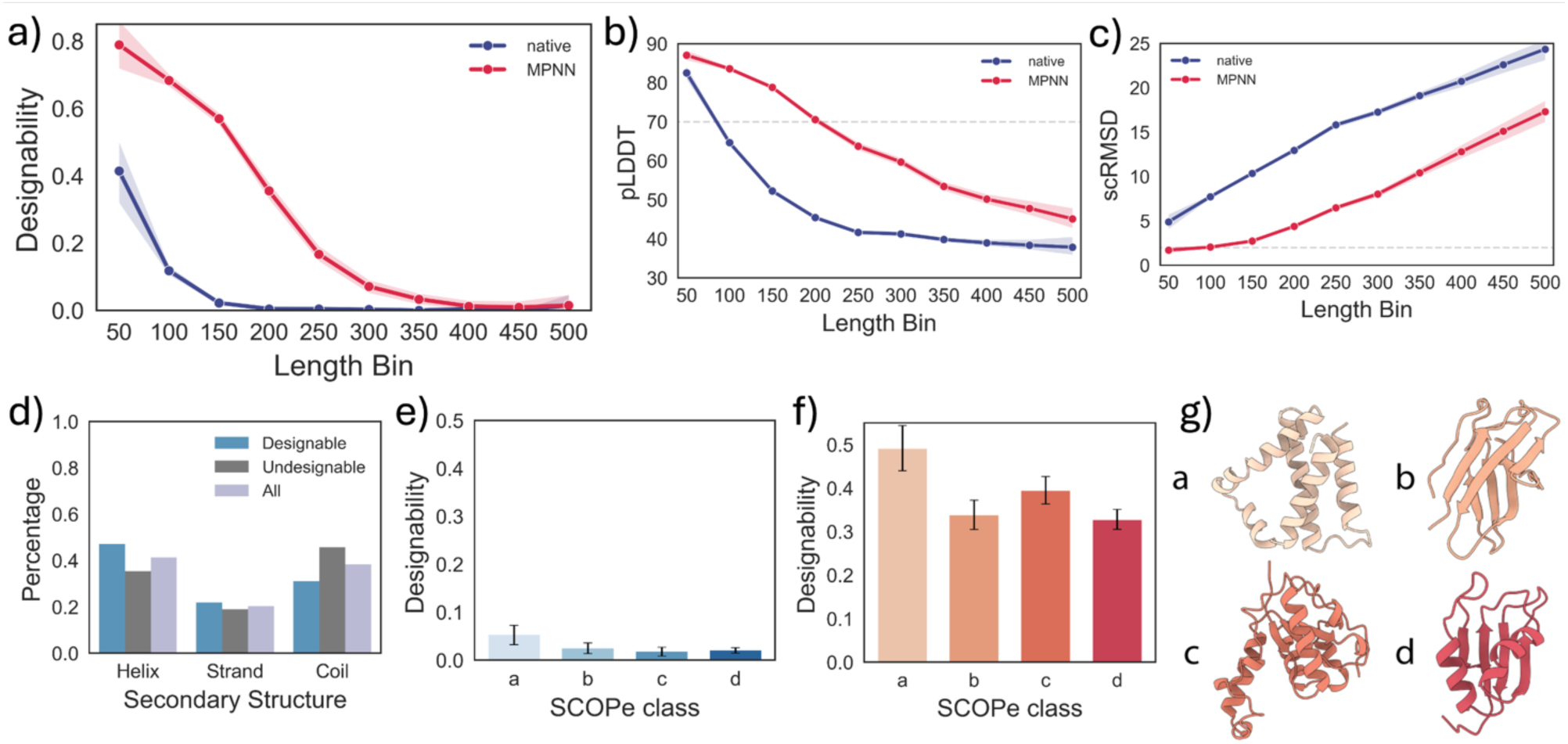
Structural characteristics affecting designability of the SCOPe dataset. a) Designability of proteins from SCOPe as a function of their length. Scatter points report mean values for a given length bin. b-c) pLDDT and scRMSD values of the proteins from a). Designability thresholds for pLDDT ≥ 70 and scRMSD ≤ 2.0 Å are depicted as grey dashed lines. d) Percentage of helices, strands, and coils for structures from natural SCOPe sequences. d) Percentage of designable sequences from SCOPe domains and f) their pMPNN designs ordered by SCOPe class. For each class, a uniform length distribution between 60 and 300 residues of pMPNN-designed sequences was randomly selected from the initial SCOPe dataset. g) Visualization of representative structures for SCOPe classes. a: all-α, b: all-β, c: α/β (mixed), d: α+β (segregated).

When we scrutinized designable and undesignable SCOPe proteins for structural characteristics encoded by their natural sequences, we identified an overrepresentation of α-helices compared to β-strands or coils (Fig. 4d) in all designable sequences. To test if this bias persists across SCOPe classes independent of sequence length, we selected a uniform length distribution between 60 and 300 residues for each SCOPe class. Investigating the designability of both native sequences (Fig. 4e) and the lowest-scRMSD pMPNN-designs (Fig. 4f), we found that class a representing α-helical domains systematically achieved higher fractions of designable sequences than other classes.

These results reveal the conundrum protein designers face when using AF2 ss as an oracle: if it can fold a significant fraction of sequences in a set, it can indeed be leveraged to enrich the fraction of foldable designs in candidate sets. However, for sequences which do not exhibit as tight a sequence-to-structure mapping as pMPNN designs, and which encode even medium-sized or mixed secondary structure proteins, the inability to accurately predict any structures severely limits its use as an oracle.

### 2.4 Refolding-based designability assessment is sensitive to disordered and flexible regions

Observing that both sequence and structure properties influence designability distribution and thus the reliability of refolding metrics as predictors of experimental success, prompted us to examine other factors that could affect performance. We suspected that one clue might lie in the pronounced designability gap for AlphaFold Database (AFDB) structures. As observed by Lin et al.,^43^ pMPNN-redesigned AFDB structures show only about half the designability of designs generated from natural structures taken from PDB. The authors hypothesized that the observed discrepancy might arise from atomic inaccuracies in AFDB models or limited generalization of sequence design models trained solely on PDB data.^43^ Similarly, designability-based performance drops were also observed when using high-quality AF2 predictions for training backbone design models.^34^ These findings prompted us to investigate particularities in AFDB structures that may render refolding-based designability assessment unreliable.

We first hypothesized that the lower average designability of AFDB structures might stem from out-of-distribution folds, i.e. domains with no sequence and structure similarity to experimentally resolved structures. These so-called “dark clusters” might be less amenable to inverse folding, driving designability down.^44^ We thus compiled datasets of high-pLDDT (>80) AFDB structures of either “light” or “dark” FoldSeek cluster representatives and included a set of SCOPe PDB domains as a baseline. Each set consisted of 1000 random proteins selected to represent identical length distributions. We then designed 8 sequences per structure with pMPNN and, for comparability with Lin et al.,^43^ predicted structures with ESMfold, before computing designability for the lowest-scRMSD design. While we were able to reproduce the previously observed designability gap, the designability metrics were similarly lowered for dark and light cluster representatives (Fig. 5a).

**Figure 5.**
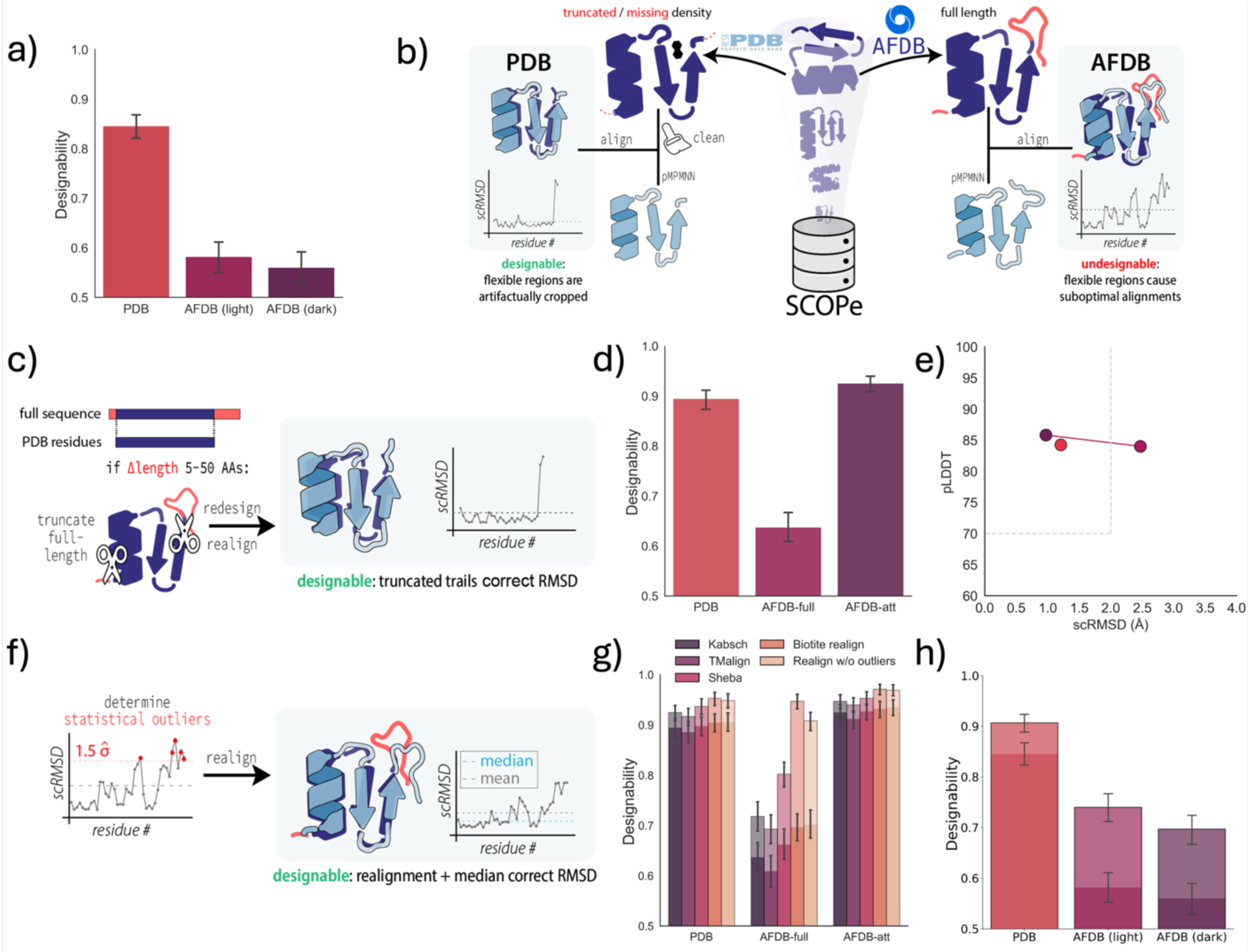
**a)** Designability of the best-ranked pMPNN designs of the PDB compared to the AFDB structures from either light or dark FoldSeek cluster representatives of AFDB. b) Schematic illustration of the superposition of ESMfold-predicted structures of pMPNN designs onto PDB and full-length AFDB structures, where disordered regions cause misalignment and inflated scRMSD. c) Procedure for the attenuation experiment (AFDB-att): structures with a 5-50 amino acid length difference between PDB and AFDB are truncated before alignment. d) Designability of the PDB, full-length AFDB (AFDB-full), and AFDB attenuated to PDB length (AFDB-att) designs. e) Mean pLDDT and RMSD values for PDB, AFDB-full, and AFDB-att. f) Correcting misalignments due to disordered regions: outliers in the per-residue scRMSD are excluded during realignment, and the median scRMSD over all residues is taken. g) The effects of different structure alignment methods are compared using either the mean (no outline) or median (black outline) scRMSD. h) Designability data shown in a) (no outline) corrected with the median realigned scRMSD (black outline).

Although these results led us to exclude dark clusters as the primary cause of reduced designability in AFDB-based designs, they prompted us to examine structural differences between the datasets. Notably, secondary structure recurrently impacted our analyses: the high-pLDDT dark cluster representatives were enriched in α-helices (Fig. S9), which potentially offset a more challenging structure prediction of out-of-distribution folds. Similarly, among designable sequences in our SCOPe set, helices were over, and coils underrepresented (Fig. 4d). Yet, when comparing overall secondary structure content, we found that the AFDB set is not “disadvantaged” by lower helix or higher coil content (Fig. S9). We then recognized a key distinction between the two databases, however: PDB structures frequently lack electron density for flexible loops and termini, and deposited structures are often deliberate truncations of disordered or insoluble domains (Fig. 5b). Conversely, AFDB entries are predictions of the full-length sequence and often span disordered regions within otherwise confidently predicted models. Indeed, low pLDDT intrinsically disordered regions constitute a substantial proportion of AFDB models,^45,46,47^ and differences in architecture and training across folding models may cause different models to not converge on the same conformation. As a consequence, when computing designability, scRMSD of ESMfold (or any other folding model) predictions to the target AFDB structures could become artificially high, even though the structured part of the protein is predicted correctly and results in an optimal alignment.

To test this hypothesis, we selected 1000 random SCOPe proteins in which the corresponding AFDB entries differed by 5-50 residues at the termini, and truncated the full-length AF2 predictions to match the PDB-deposited length. In this way, we obtained three datasets for the same selection of proteins: PDB, full-length AFDB structures (AFDB-full), and attenuated AFDB structures (AFDB-att, Fig. 5c). We again applied the refolding pipeline to these datasets by generating 8 pMPNN designs per structure, predicting structures with ESMfold, and computing designability for the lowest-scRMSD design. In contrast to full-length models, AFDB-att structures exhibited the same high designability as observed for the PDB (Fig. 5d). When disentangling the two metrics, we found that low designability of AFDB-full is not driven by lower pLDDTs, but substantially higher scRMSD values (Fig. 5e), corroborating our hypothesis. As scRMSD depends on structural alignment, this result may be explained by two effects: either the flexible termini in full-length predictions inflate the mean scRMSD, or truncation improved the global alignment by preventing flexible regions from biasing the fit (Fig. 5b).

To address this question without having to rely on experimental structures to explicitly identify flexible regions as we did in the attenuation approach, we implemented an adapted pipeline for correcting the designability metrics: to avoid the effects of structural misalignment due to unstructured parts, we realigned the structures under exclusion of all outliers in the per-residue scRMSD after an initial alignment (Fig. 5f). Moreover, we took the median instead of the mean scRMSD, to prevent an inflated average after realignment. In addition to this simple custom approach, we also compared established tools that could improve the outcome of using the default Kabsch alignment method: biotite’s “superimpose_without_outliers,”^48^ which also removes statistical outliers but iterates until convergence; Sheba,^49^ which considers residue environments in 1D and iteratively identifies matching pairs in 3D; and TMalign, which optimizes the Kabsch rotation matrix using the global TM-score.^50^ While TMalign and Sheba did not rescue the AFDB-full set to the designability of the PDB or AFDB-att sets, both approaches of realigning after outlier rejection resulted in a full rescue when combined with the median scRMSD (Fig. 5g). This outcome indicates that the presence of flexible outlier regions can indeed result in significant global misalignments, and that a simple RMSD-based Kabsch alignment after a single round of identifying outliers can mitigate the issue.

As this analysis was performed on a set of structures preselected to contain flexible tails, we next sought to validate the observed rescue in an unbiased set. To that end, we used the previously collected light and dark cluster AFDB representatives (Fig. 5a) and calculated the median scRMSDs after single outlier-rejected re-alignment. Again, the procedure resulted in a marked increase in designability for both light and dark cluster representatives, indicating that flexible termini in the AFDB contribute to a significant decrease in designability of the AFDB compared to the PDB (Fig. 5h).

In summary, we conclude that flexibility is an additional factor that can reduce the utility of refolding metrics, besides sequence length and secondary structure content. Our analysis also suggests that the previously observed reduced designability of the AFDB likely arises from misalignments between target and designed structures driven by flexible or disordered regions (Fig. 5d, h), and that this issue is remedied by a simple alignment correction step.

### 2.5 Designability assessment is only moderately affected by folding and sequence design settings

Having characterized the limitations of the refolding pipeline and designability metrics, we finally asked whether straightforward workflow adjustments could mitigate these. Using the so-far default structure prediction and sequence design settings as a baseline, we varied the number of AF2 recycles and the number of sequences generated with pMPNN, two key parameters likely to affect refolding metrics and commonly adjusted in protein design studies.^11,16,24^

We randomly selected 500 proteins from our SCOPe dataset and first refolded either their natural sequences or the lowest (out of 8) scRMSD-ranked pMPNN designs using AF2 ss with varying recycles. In contrast to natural sequences, where designability remained very low regardless of the number of recycles, AF2 ss predicted designability threshold-passing structures with high confidence for 20% of pMPNN designs even without recycling. Although the highest designability of 50% was only reached after 48 recycles (at a runtime of about 20 times more compute than 0 recycles), saturation (>90% of maximum) occurred already at 12 recycles. Interestingly, we saw that picking one random from the resulting five AF2 models on average causes choosing a substantially less confident prediction, but that this effect was much more pronounced for pMPNN designs than for native sequences (Fig. 6b). Curiously, when checking model-specific effects by analyzing how often each of the five resulting AF2 models produced the highest pLDDT, we found model 2 to most frequently perform best (Fig. S10a), and model 1 most often worst. In absolute pLDDT units, the difference between a random and the best-performing model was very similar across different models, however (Fig. S10b).

**Figure 6.**
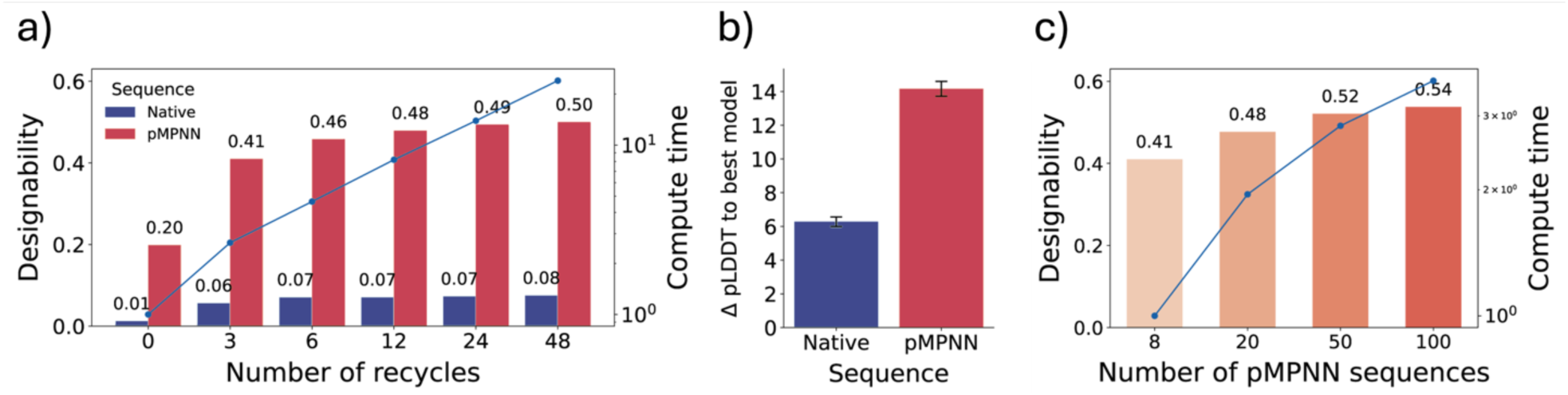
Effect of different AF2 ss structure prediction and pMPNN sequence design settings on the designability of 500 randomly selected SCOPe proteins. a) Designability of natural and lowest-scRMSD pMPNN sequence as a function of the number of recycles. b) pLDDT difference between randomly choosing a single AF2 model (out of five) and the model yielding the highest-confidence prediction for the proteins in a). c) Designability of the same set as in a) with 3 recycles, but with varying number of pMPNN sequences.

We next varied the number of sequences designed with pMPNN from the default 8 up to 100, keeping the default sampling temperature T = 0.1 and the model’s training noise level of 0.2 Å, as shown to be optimal for the highest *in silico* success rate.^16^ Notably, sampling more sequences with pMPNN at a fixed number of three recycles increased the fraction of designable samples to a higher extent than increasing recycles did (Fig. 6c). In our benchmark, 100 instead of 8 sequence designs took only about four times longer.

Together, these experiments show that the low fraction of designable native sequences can only be remedied to a limited extent by increasing recycles as well as predicting and selecting from all five models. Conversely, designability with pMPNN sequences, which are already easier to fold for AF2 in general, can additionally be boosted by simply designing more sequences.

## 3 Discussion

With the growing accessibility and ease of use of protein design tools, the field is rapidly expanding and attracting researchers from diverse areas of the biosciences. Refolding-based designability assessment has become standard practice in nearly all design studies. Here, we systematically evaluated this approach and highlighted several limitations that may not be immediately evident, particularly to newcomers in the field.

We first show that the inclusion of evolutionary information compromises the reliability of both AlphaFold2 and ESMfold in evaluating sequence-structure compatibility. This is evident from AF2’s poor ability to discriminate between nonsensically mutated variants and their natural or well-designed counterparts. These findings extend the widely held view that structure prediction models cannot reliably infer the effects of single mutations on stability or fold.^51,52^ One can speculate that the underlying causes might be similar: the dominance of strong sequence-to-structure signals from wild-type residues, the lack of explicit physics-informed losses incentivizing folding models to predict mutation effects, and the absence of non- or misfolded proteins in the training data.

Crucially, the use of MSAs markedly diminishes the predictive power of pLDDT and scRMSD for experimental success, as shown by our analysis of literature-obtained sequences with known experimental outcomes. Although ESMfold does not explicitly use MSAs, its performance typically falls between that of AF ss and AF MSA. Since ColabFold—one of the most widely used prediction platforms—computes MSAs by default, our findings call for caution when redesigning natural proteins, where refolding metrics are most easily confounded. The continued use of MSA mode in numerous design studies^11,29,30,31,32^ and meta-analyses^41^ highlights the need for a critical reassessment of current best practices.

Our results also support the general notion that confidence metrics from AF2 single sequence mode, and to a lesser extent by ESMfold, are correlated with experimental success. However, AF2 ss fails to predict correct structures of many natural sequences, rendering it unreliable for assessing designability when the sequence identity of a design to a natural sequence is high, even with many recycles. AF2 ss performance also depends on sequence length and structural composition, with all-α proteins folding most successfully. Notably, pMPNN-designed sequences tend to receive high pLDDT and low scRMSD values, underscoring a key limitation of refolding: structure prediction models are not true oracles of stability. They capture protein energetics only implicitly and may assign high confidence to sequences that are structurally plausible *in silico*, but biophysically unstable or non-functional in reality. This limitation is also relevant for model development: efforts such as training sequence design models by rewarding designability risks further inflating false positives and could ultimately obsolete the metric for oracle use.^23^

Finally, we identify a broader limitation in the current designability definition—its dependence on full-body superposition when calculating scRMSD. This superposition is highly sensitive to flexible or disordered regions that lack a stable conformation and may adopt different states under different folding runs and models. Such regions can inflate scRMSD values beyond designability thresholds even when the structured part is correctly predicted and aligned. This straightforward explanation of the observed designability discrepancies between experimental and predicted structures allowed us to propose attenuation or a simple outlier rejection and realignment procedure to alleviate the issue. This insight can help complementing approaches aiming to “debias” the AFDB,^53^ and our findings may help explain decreased performance of classifiers^54,55^ and generative models^56^ when switching between experimental and predicted structures. Our results also align with previous observations that higher structural flexibility is correlated with increased scRMSD in small monomeric SCOPe proteins.^57^ Taken together, our findings indicate that the prevailing designability definition inherently favors compact, rigid structures and underrepresents flexible or disordered regions. As the protein design field increasingly seeks to encode flexibility, we hope that our findings underscore the need for greater awareness of this bias and for developing more adequate metrics in the future.

## 4 Methods

### 4.1 Self-consistency pipeline

The goal of the self-consistency (refolding pipeline) is to *in silico* assess whether a designed sequence encodes a target protein structure, i.e. estimate the probability of a structure given a sequence. If a refolded natural or designed sequence has a pLDDT higher and a self-consistency root mean square deviation (scRMSD) to the target structure lower than a certain pre-defined threshold, the sequence is called *designable*. We depict a schematic visualization of the refolding pipeline in Fig. 1.

#### Designability computation

Designability evaluation based on refolding metrics is well-established in the literature,^8,21,26,35^ and relies on both pLDDT and scRMSD. Unless stated otherwise, scRMSD is computed by aligning the C*α* coordinates of the predicted model to the target structure using an adapted version of the Kabsch algorithm, available at: https://github.com/nghiaho12/rigid_transform_3D. We report pLDDT as the average per-residue value over the full protein length. Unless explicitly stated otherwise, we consider a sequence designable if it achieves pLDDT ≥ 70 and scRMSD ≤2.0 Å.

#### Sequence design with ProteinMPNN

Whenever we designed sequences with ProteinMPNN (pMPNN), unless stated otherwise, we used the default inference settings with model v_48_020: sampling temperature *T* = 0.1, backbone noise level during training =0.2 Å, and 8 sequences per design. After refolding, the pMPNN sequence with the lowest RMSD to the target backbone was used for further evaluation.

### 4.2 Structure prediction

In our experiments, we predicted structures of a given set of sequences using four different approaches: with AlphaFold2 (AF2) using either (1) single sequence mode with no MSA (AF2 ss), (2) using an MSA (AF2 MSA), (3) using an MSA where the columns of mutated residues are masked (AF2 MSA mask) or (4) using ESMfold.

The full MSAs were generated with MMseqs2 using the colab_search implementation provided by ColabFold 1.5.2, with default settings. No templates were provided, and the databases uniref30_2202_db and colabfold_envdb_202108_db were used for the DB1 and DB3 parameters, respectively. For the AF2 MSAs with masks, we removed native sequence context from the MSA by masking entire columns with the token X, and performed structure predictions using the same settings as for the unmasked AF2 MSAs.

All AF2 predictions were performed with a locally installed ColabFold 1.5.2 version with 3 numbers of recycles and 5 different models if not explicitly stated otherwise. In all cases, the highest-ranked plDDT model was used. We provide no templates for the AF2 predictions and do not relax them. For AF2 single sequence mode predictions, we provide no MSA (msa_mode = single_sequence) to the model and keep other settings as described above. For structures predicted with ESMfold, we used default settings with 3 numbers of recycles and a single model output. We ran all predictions on an NVIDIA A100 (40 GB) GPU. Reported inference times in Fig. 6 were also computed with ColabFold in batch mode.

### 4.3 SCOPe dataset

As an initial dataset, we used the SCOPe dataset^58^ split by 40% sequence identity (astralscopedom-seqres-gd-sel-gs-bib-40-2.08.fa, provided at https://scop.berkeley.edu) to ensure minimal sequence homology and structural redundancy between proteins. We further subset the SCOPe dataset to include single-domain proteins with lengths between 50 and 512 residues. For additional details about dataset subsets created for specific experiments, we refer the reader to the corresponding Methods sections described below.

### 4.4 Mutagenesis of natural proteins

#### Selection of proteins

We randomly selected 50 small, distinct protein domains from the SCOPe dataset introduced in Methods 4.3 that are classified as designable if predicted with AF2 single sequence mode. To avoid the selection being dominated by small helical proteins, we ensured that each protein included had a length greater than 60 residues and a helical content between 30% and 60%. For all sequences, abundant evolutionary information in the form of an MSA is present (Fig. S2, SCOPe wt). Mutagenesis of natural sequences. Starting from native sequences, we gradually mutated from 10% to 60% of residues to random natural amino acid tokens. For each fraction of mutated residues, we generated 64 sequences to reduce the impact of sequences that may contain less disruptive mutations by chance. In total, this dataset contains 19200 protein sequences. Full-length MSAs and masked column MSAs were generated for each sequence as described in Methods 4.2. In this experiment, we computed designability as a fraction of predicted models passing the criteria defined as: scRMSD ≤ 2.0 Å and pLDDT ≥ (70, 80, 85). We vary the pLDDT threshold since pLDDT shows variability across different protein design studies, as explained in the introduction.

#### Mutagenesis of ProteinMPNN designs

To further evaluate the sensitivity of designability to random mutations in the context of pMPNN-designed sequences, we repeated the experiment, mutating the pMPNN designs instead of natural ones. For each of the 50 proteins, we generated 8 pMPNN sequences and refolded them using the AF2 ss. The pMPNN-designed sequence with the lowest scRMSD to the target structure was used as input for the same mutagenesis experiment as described for the native sequences.

#### Biophysical metrics

To demonstrate the unphysical nature of the mutated sequences, the energy and aggregation propensity were calculated for all mutated sequences that passed the designability thresholds in AF2 MSA mode. The Rosetta energy and spatial aggregation propensity (SAP) were calculated using the PyRosetta,^39^ using the predicted models generated by AF2 MSA as poses. The energy of the poses was calculated using the default energy function (ref2015),^4^ with the output reported in Rosetta Energy Units (REU). The SAP scores were calculated using the SapScoreMetric() class in the pyrosetta.rosetta.core.pack.guidance_scoreterms.SAP module.

### 4.5 Effect of evolutionary information on refolding metrics as proxies of experimental success

#### SoluProt Dataset

Entries from the SoluProt database were matched with their PDB IDs, and all entries for which an experimental structure was available were selected, resulting in 925 entries of 74 insoluble and 851 soluble proteins. To control for length, the largest possible number of soluble entries were selected that matched the same length distribution as the 74 insoluble entries using 10 bins, resulting in a set containing 74 insoluble and 220 soluble entries (Fig. S3). Structures of all entries were predicted using AF2 single sequence mode, AF2 MSA, and ESMFold as reported in Methods 4.2. Significance was calculated with Fisher’s exact test (*p* < 0.05). To control for any biases introduced in the above selections, a further 1000 entries were randomly selected from the SoluProt database and ran in the same manner described above, this time only calculating pLDDT as most entries do not have an experimental ground truth structure available (Fig S4).

#### Dataset of experimentally tested designs

To compare differences in the refolding pipeline to experimental outcomes, we obtained experimentally verified sequences from 7 different publications: (1) A set of 42 water-soluble *de novo* β-barrels generated using ab initio energy calculations obtained from Dou et al.^59^ (2) a set of 81 transmembrane *de novo* β-barrels generated using ab initio energy calculations obtained from Vorobieva et al.^60^ (3). A set of 85 membrane proteins solubilized with AF2seq and pMPNN obtained from Goverde et al.^25^ (4). A set of 70 *de novo* designs generated by the Foldit community and experimentally verified by Koepnick et al.^61^ (5). A set of 87 *de novo* P-loop and Rossman fold-like proteins generated using ab initio energy calculations obtained from Koga et al.^62^ (6) a set of 40 novel α-helical proteins generated using assembled helix fragments by Sakuma et al.^63^ (7) a set of 60 novel α/β-fold proteins generated using fragment assembly simulations by Minami et al.^64^

#### Evaluation

Structures of all sequences were predicted using AF2 single sequence mode, AF2 MSA, and ESMFold as reported in Methods 4.2. Some of the data, such as the β-barrels reported by Hermosilla et al., required 48 recycles to fold properly. For completeness, we report metrics for both the default 3 and adjusted 48 recycles. The ROC curves independent of thresholds for both pLDDT and RMSD are reported. Median per-residue Neff scores were calculated as reported in Jumper et al.^1^

Performance of different threshold regimes for either precision (prioritizing high success rates) or F1 score (balancing high success rates with identifying sufficient hits) was investigated by varying the pLDDT threshold between 50 and 100 in steps of 5, and the scRMSD threshold between 0 and 5 in steps of 0.5Å. To guarantee well folded structures for all datasets, the metrics using 48 recycles were used. Optimized thresholds were selected based on the maximum precision or F1 scores. When multiple threshold combinations resulted in the same maximum precision or F1 score, the most stringent set of thresholds was chosen to facilitate closer clustering of the thresholds where possible. Confusion matrices were calculated for the optimized F1 thresholds to demonstrate the increase in the true positive rate between AF2 ss and AF2 MSA, and heatmaps were plotted to indicate the difference in performance between AF2 ss and AF2 MSA under all investigated threshold regimes.

### 4.6 Limitations of designability assessment with AF2 ss

#### Designability of natural proteins

In this experiment, we ran the refolding pipeline on the whole SCOPe dataset described in Methods 4.3. Besides filtering for length filter between 50 and 512 residues, we filtered the SCOPe to contain only complete, single-domain proteins, and further excluded small and membrane proteins from the dataset (classes f and g). Finally, we excluded two abundant repetitive SCOPe folds a.118 (alpha-alpha superhelix) and b.69 (7-bladed beta-propeller) which would otherwise result in artifacts in the designability/length distribution. These refinements resulted in a subset of 9916 of the original 15177 proteins from the SCOPe dataset. We predict the structures of natural or lowest scRMSD-ranked pMPNN designed sequences using AF2 single sequence mode with designability defined as pLDDT ≥ 70 and RMSD ≤ 2.0 Å, as described in Methods 4.1. The effect of secondary structure was evaluated using the pyDSSP implementation of the DSSP algorithm (https://github.com/ShintaroMinami/PyDSSP).

To evaluate the effect of SCOPe class on designability of pMPNN sequences, we control for the length effect by selecting predicted structures of lowest scRMSD-ranked pMPNN-designed sequences with a uniform length distribution between 60 and 300 residues for SCOPe classes a (all-alpha), b (all-beta), c (a/b proteins), and d (alpha+beta). For each class, the largest number of proteins were selected that could still guarantee a flat distribution within the selected length range, using 15 bins. This resulted in flat distributions with 126 proteins for class a, 252 for class b, 252 for class c, and 462 for class d respectively.

### 4.7 Effect of structure prediction and ProteinMPNN settings on designability computed with AF2 ss

From the SCOPe dataset described in Methods 4.3, we randomly selected 500 proteins, verifying that the length distribution was representative of the total SCOPe distribution. This set was used to conduct experiments with a varying number of AF2 recycles (0-48) and pMPNN sequences (8-100). We predict the structures of natural or pMPNN-designed sequences using AF2 single sequence mode with designability defined as pLDDT ≥ 70 and scRMSD ≤ 2.0 Å, as described in Methods 4.1.

### 4.8 Designability of the AlphaFold Database (AFDB)

#### Dark clusters in the AlphaFold Database (AFDB)

Barrio-Hernández et al.^44^ provides a FoldSeek-clustered version of the AlphaFold Database, in which clusters are classified as *dark* if none of their members can be annotated with PFAM or TIGRFAM20 domains in the UniProt/TrEMBL or SwissProt databases. From the cluster overview dataset available at https://afdb-cluster.steineggerlab.workers.dev/, to align our data with the observation made by Lin et al.,^43^ we randomly selected 1000 dark cluster representatives that fulfilled the following criteria: each protein belonged to a cluster with at least 10 members and had a representative structure with pLDDT ≥80. This set represents protein samples that are distinct from any proteins found in PDB. To compare, 1000 non-dark cluster representatives and 1000 SCOPe domains with identical length distributions were chosen using the same criteria, and are referred to in the text as the “light” and “PDB” sets respectively. To assess the effect of flexible regions on the AFDB, we use both the light and dark AFDB sets as representatives. Although this risks over-representing dark clusters, our result shows these two datasets exhibit similar designability.

#### AlphaFold database attenuation

For the SCOPe dataset described in Methods 4.3, we obtained the mappings of each protein to its corresponding UniProt entry from the Structure Integration with Function, Taxonomy, and Sequence (pdb_chain_scop_uniprot.tsv, provided at https://www.ebi.ac.uk/pdbe/docs/ sifts/quick.html) and downloaded the AF2 predictions from the AlphaFold Database (AFDB). Sequences encoding the AFDB models were extracted, and 1000 samples were randomly selected where the length difference between the SCOPe and AFDB sequences for the same UniProt mapping was greater than 5 but less than 50 residues. We attenuated the structures of AFDB models to the PDB-deposited length for subsequent analyses, as described in Section 3.5. Sequences were designed for the PDB, full-sequence AFDB, and attenuated AFDB structures using pMPNN analogously to Methods 4.1. Structure predictions were performed with ESMfold using default settings.

#### scRMSD corrections

To correct misalignment and inflated scRMSD due to flexible N- and C-terminal regions, we realigned the light and dark AFDB sets and the corresponding PDB set with different methods. We tested realignment with TMalign,^50^ Sheba, ^49^ and the “superimpose_without_outliers” function implemented in Biotite.^48^ As residues can be excluded from the scRMSD during the sequence alignment when using TMalign and Sheba, only the structural alignment functionality is used, and the final scRMSD is calculated using np.linalg.norm. The outlier-excluded realigned scRMSD is calculated using the following steps. (1) per-residue RMSD is calculated using the default Kabsch algorithm. (2) outliers among the per-residue RMSD values are selected if the per-residue RMSD is larger than the median RMSD + 1.5x the Median Absolute Deviation (MAD). (3) The scRMSD is realigned while masking the selected outlier residues. (4) Either the mean or median scRMSD of the full structure is reported.

## Acknowledgements

M.J.L.J. Fürst gratefully acknowledges funding from the Netherlands Organization for Scientific Research NWO (VI.Veni.212.263). This work was supported by the European Union through COST Action CA21162 (Establishing a Pan-European Network on Computational Redesign of Enzymes) and used the Dutch national e-infrastructure with the support of the SURF Cooperative using grant no. EINF-4326. We thank Casper Goverde for providing experimental data for the solubilized membrane protein analogues. We also thank the Center for Information Technology of the University of Groningen for providing access to the Hábrók high-performance computing cluster, and the National Academic Infrastructure for Supercomputing in Sweden (NAISS), partially funded by the Swedish Research Council through grant agreement no. 2021-29 for awarding this project access to the Berzelius resource provided by the Knut and Alice Wallenberg Foundation at the National Supercomputer Centre.

## Competing Interests

The authors declare no competing interests.

## Data and Code availability

The source code and data for all of the figures are available at https://github.com/kt-korbeld/Limitations-refolding-pipeline-data upon acceptance of the article.

## Notes

https://github.com/kt-korbeld/Limitations-refolding-pipeline-data

### Competing Interest Statement

The authors have declared no competing interest.

