## Supplementary Figures for "Limitations of the refolding pipeline for *de novo* protein design"

[‡]Equal contribution

### Figure S1

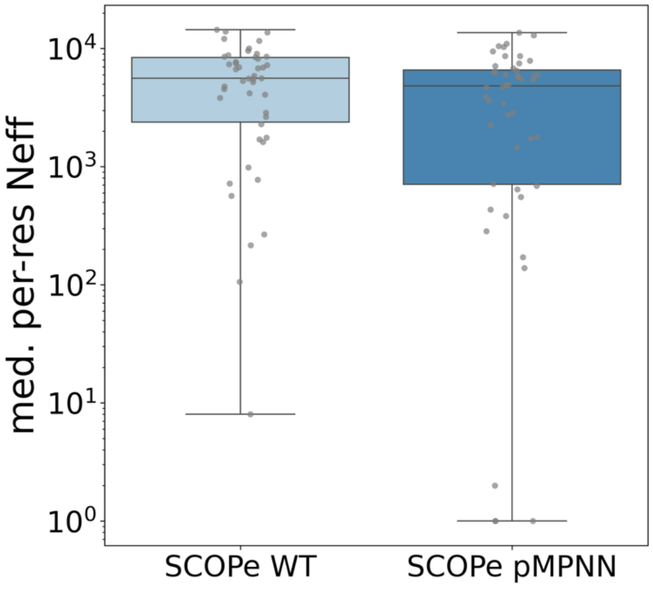

**Figure S1.** MSA depth given as the median per-residue $N_{eff}$ computed for the dataset used in Fig. 1 (50 small SCOPe proteins and their best pMPNN redesigns).

### Figure S2

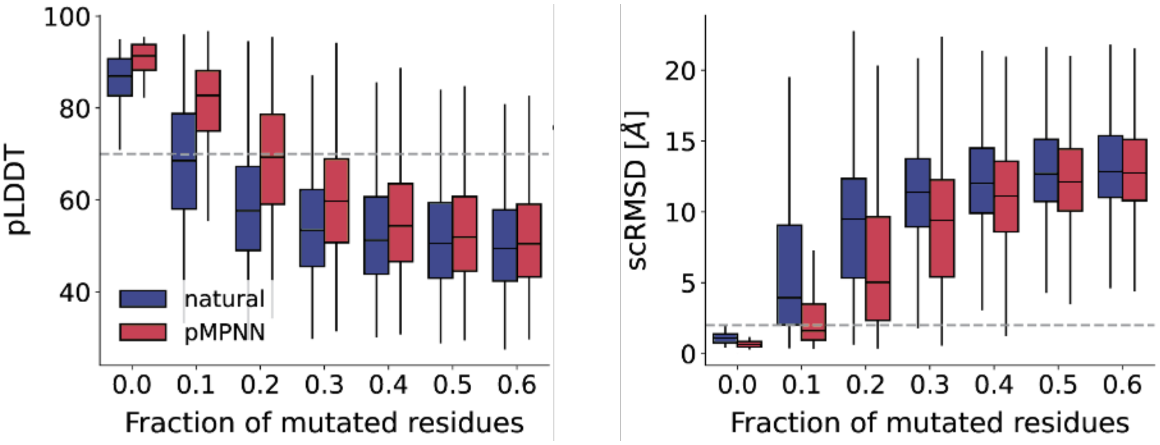

**Figure S2**. pLDDT and scRMSD values for mutated sequences derived either from natural or pMPNN designed sequences for 50 small SCOPe proteins predicted with AF2 ss for the indicated percentage of mutated residues.

### Figure S3

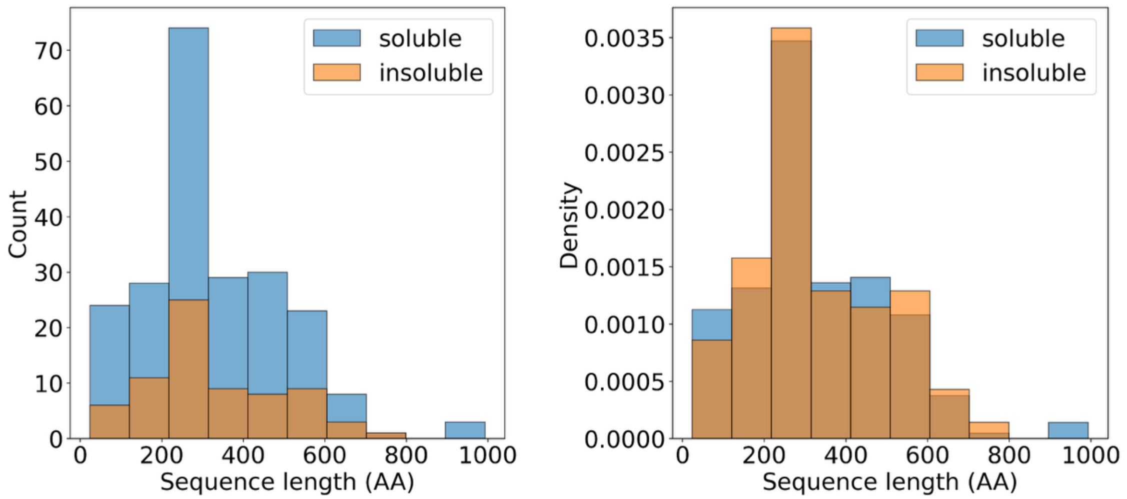

**Figure S3.** Length distributions of the selected structures for the SoluProt dataset, shown as absolute count (left) or density (right). As soluble entries were more abundant than insoluble, the length distribution of the soluble sequences was matched to the length distribution of the insoluble sequences, resulting in 74 insoluble entries and 220 soluble entries.

### Figure S4

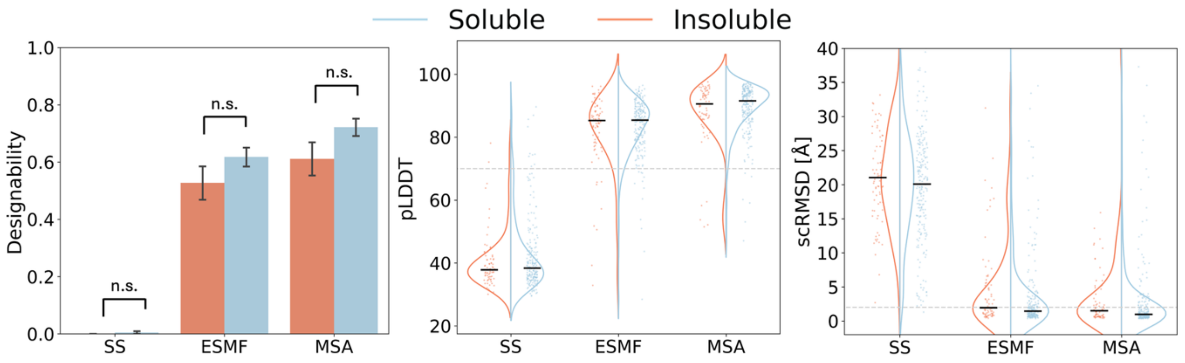

**Figure S4.** Designability, pLDDT, and scRMSD metrics for 294 proteins from the SoluProt dataset (Fig. S3) showing that AF2 and ESMfold are unable to distinguish natural proteins that lead to soluble expression in heterologous overexpression from those that do not, regardless of whether an MSA is provided as input or not.

### Figure S5

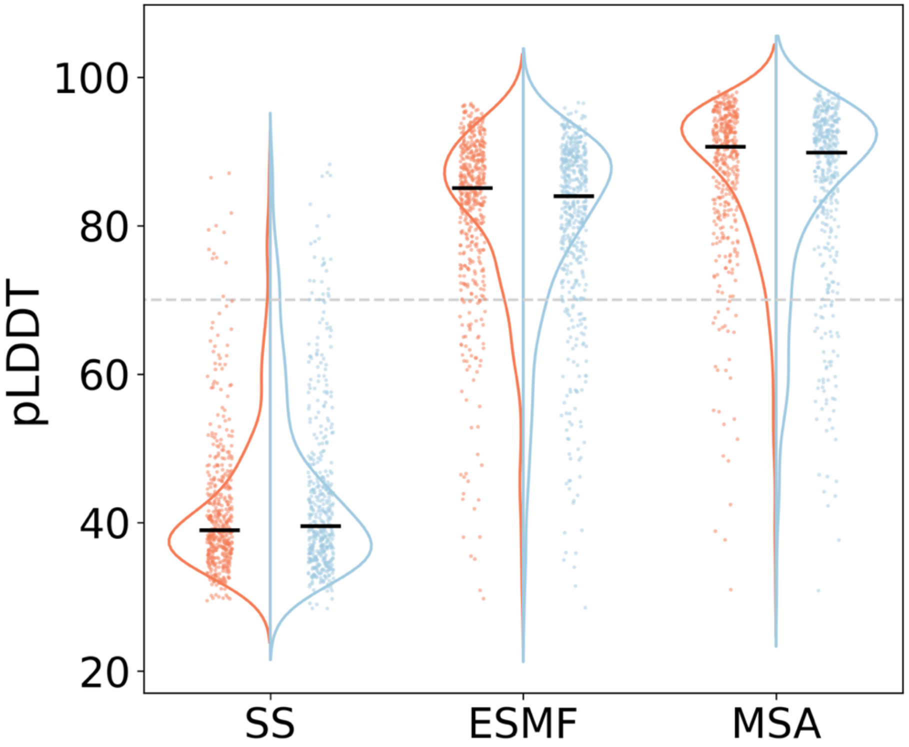

**Figure S5.** pLDDT for 1000 random proteins from the SoluProt dataset showing that AF2 and ESMfold are unable to distinguish natural proteins that lead to soluble expression in heterologous overexpression from those that do not, regardless of whether an MSA is provided as input or not.

### Figure S6

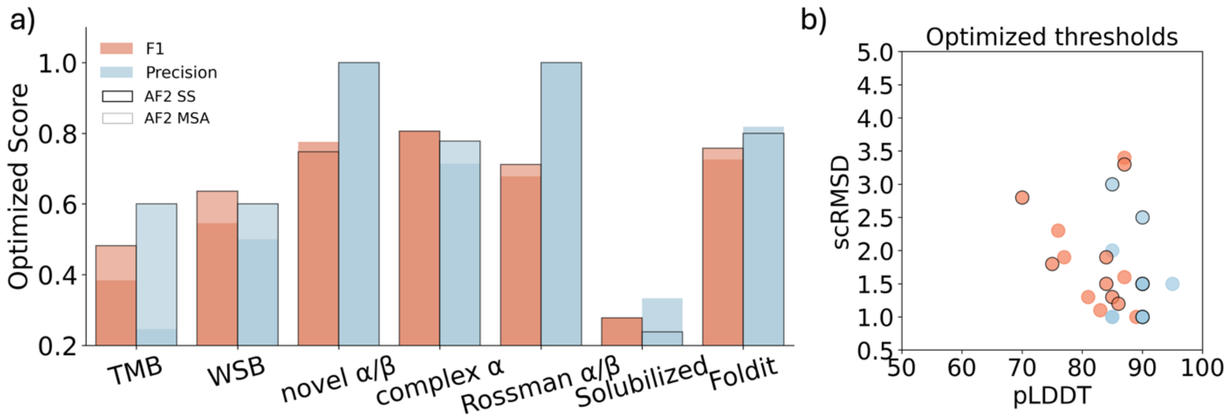

**Figure S6**. Thresholds are optimized for either precision or F1 score within a range of 50-100 for pLDDT and 0.5-5Å for scRMSD. a) the resulting optimized F1 and precision scores across each dataset. Datasets are ordered by MSA depth (see Fig. 3c). AF2 SS (bold outline) consistently outperforms AF2 MSA (no outline) in both F1 and precision for the datasets with deep MSAs. b) the resulting optimized thresholds for F1 or precision and AF2 SS or AF2 MSA show a broad distribution. The strictest thresholds were chosen when multiple optima exist.

### Figure S7

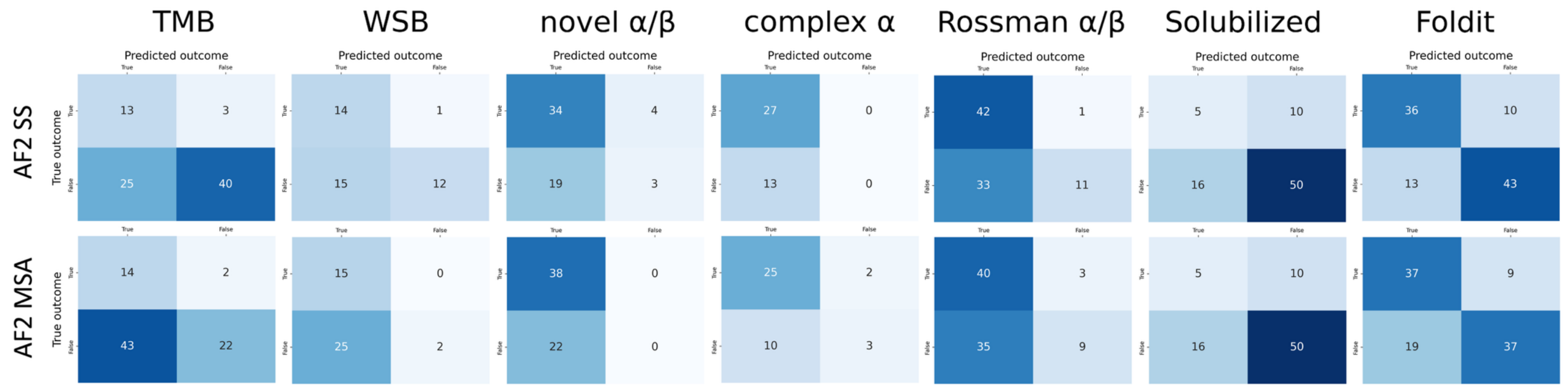

**Figure S7.** Confusion matrices for thresholds optimized using the F1 score (Fig. S6). Datasets are sorted by MSA depth from left to right (see Fig. 3c). The decreases in F1 performance are primarily driven by increases in the false positive rate when switching from AF2 SS to AF2 MSA. Datasets with deep MSAs (left) are most affected.

### Figure S8

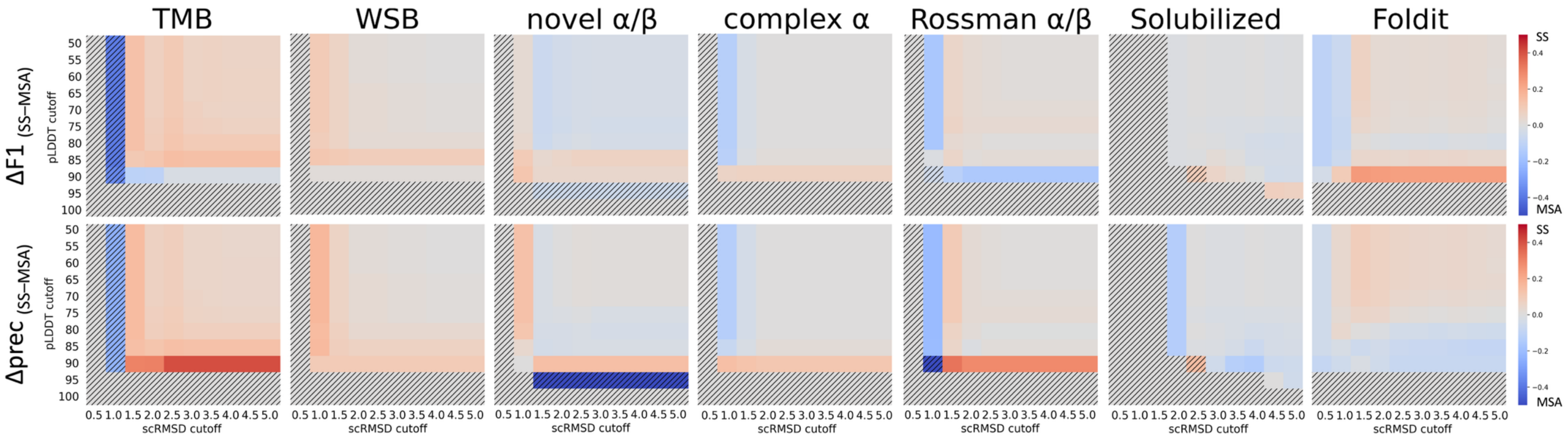

**Figure S8.** For both F1 and precision scores, the difference in performance between AF2 SS and AF2 MSA is given as a function of pLDDT (in steps of 5) and scRMSD (in steps 0.5Å) thresholds across various protein (re)design datasets from literature. Red indicates threshold regimes where AF2 SS outperforms AF2 MSA. Blue indicates threshold regimes where AF2 MSA outperforms AF2 SS. Thresholds at which recall for AF2 SS or AF2 MSA became 0 (i.e. not a single sequence was deemed designable anymore) are indicated with black stripes. Datasets are sorted by MSA depth from left to right (see Fig. 3c).

### Figure S9

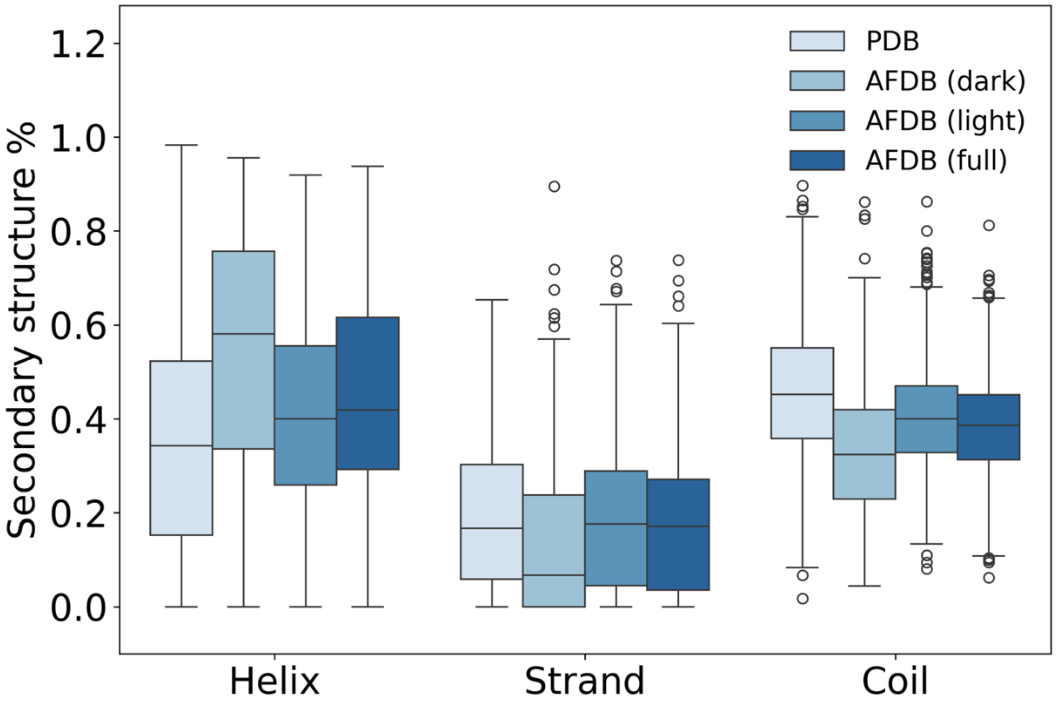

**Figure S9.** Secondary structure composition of 1000 SCOPe PDB structures (PDB), 1000 AFDB structures from dark cluster representatives (AFDB dark), 1000 AFDB structures from light cluster representatives (AFDB light), and 1000 unfiltered AFDB structure (AFDB full).

### Figure S10

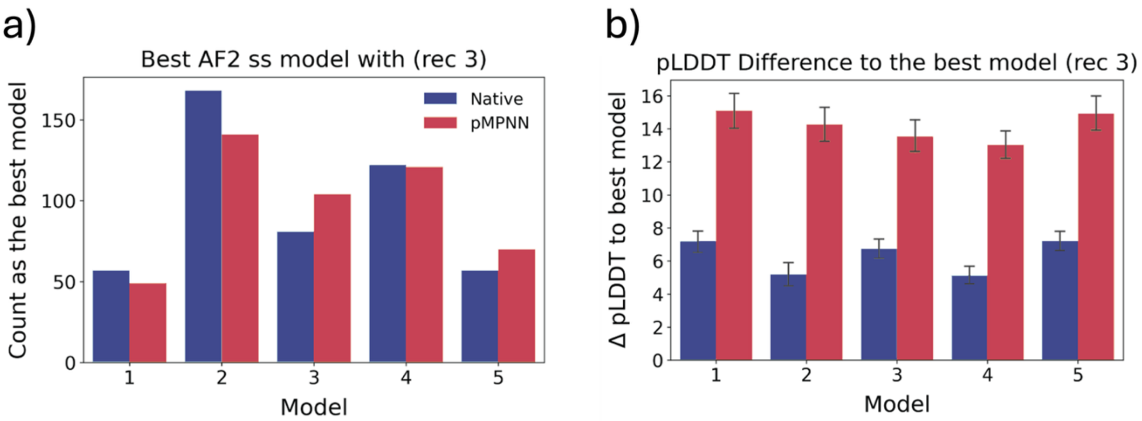

**Figure S10.** Analysis of 5 different AF2 models for 500 SCOPe proteins from Fig. 6. (a) Number of times a given AF2 model (on x-axis) yielded the best pLDDT value for a prediction in single sequence mode. (b) Difference in pLDDT to the best model prediction if a random model is taken.
